# GABA-ergic dynamics in human frontotemporal networks confirmed by pharmaco-magnetoencephalography

**DOI:** 10.1101/803924

**Authors:** Natalie E. Adams, Laura E. Hughes, Holly N. Phillips, Alexander D. Shaw, Alexander G. Murley, Thomas E. Cope, W. Richard Bevan-Jones, Luca Passamonti, James B. Rowe

## Abstract

To bridge the gap between preclinical cellular models of disease and *in vivo* imaging of human cognitive network dynamics, there is a pressing need for informative biophysical models. Here we assess dynamic causal models (DCM) of cortical network responses, inverted to magnetoencephalographic observations during an auditory oddball roving paradigm in healthy adults. This paradigm induces robust perturbations that permeate frontotemporal networks, including an evoked ‘mismatch negativity’ response and transiently induced oscillations. Here, we probe GABAergic influences of the networks using double-blind placebo-controlled randomised-crossover administration of the GABA re-uptake inhibitor, tiagabine (oral, 10mg) in healthy older adults. We demonstrate the facility of conductance-based neural mass mean-field models, incorporating local synaptic connectivity, to investigate laminar-specific and GABAergic mechanisms of the auditory response. The neuronal model accurately recapitulated the observed magnetoencephalographic data. Using parametric empirical Bayes for optimal model inversion across both sessions, we identify the effect of tiagabine on GABAergic modulation of deep pyramidal and interneuronal cell populations. Moreover, in keeping with the hierarchical coding of beliefs and sensory evidence, we found a transition of the main GABAergic drug effects from auditory cortex in standard trials to prefrontal cortex in deviant trials. The successful integration of pharmaco-magnetoencephalography with dynamic causal models of frontotemporal networks provides a potential platform on which to evaluate the effects of disease and pharmacological interventions.

**Significance Statement:** Understanding human brain function and developing new treatments require good models of brain function. We tested a detailed generative model of cortical microcircuits that accurately reproduced human magnetoencephalography, to quantify network dynamics and connectivity in frontotemporal cortex. This approach correctly identified the effect of a test drug (tiagabine) on neuronal function (GABA-ergic dynamics), opening the way for psychopharmacological studies in health and disease with the mechanistic precision afforded by generative models of the brain.

## Introduction

The development of biophysically informed models of cognition and cognitive disorders would facilitate the effective translation of the mechanisms and treatments of disease. In recent years there has been progress towards detailed generative models that replicate neurophysiological correlates of cognition based on cellular and network dynamics. Such ‘Dynamic Causal Models’ (DCM) make spatiotemporal and spectral predictions that approximate observations by functional magnetic resonance imaging or electro- and magneto-encephalography (MEG) (Moran et al., 2013). To be most useful, these models should incorporate laminar, cellular and synaptic functions (Bastos et al., 2012), and adhere to basic principles of cortical connectivity (Shipp, 2016), while also being sufficiently tractable and accurate to study human cognition.

The DCM framework developed to meet these criteria, with applications in health and neurological disorders (Kiebel et al., 2008; Stephan et al., 2008; Boly et al., 2011; Marreiros et al., 2015). DCM models draw on empirical priors for synaptic time constants and conductances, together with a mean-field forward model for each major neuronal class. For each brain region, subject and condition the models’ parameters are optimised by inversion to neurophysiological data. Although such models are supported by extensive data for face-validity (Stephan et al., 2008, 2015) and construct-validity (Razi et al., 2015), it is critical that they also achieve predictive validity (Moran et al., 2014; Gilbert and Moran, 2016; Shaw et al., 2018).

We therefore undertook DCM of human frontotemporal cortical networks during a roving auditory oddball paradigm, during which sequences of tones are presented that intermittently change in frequency. The first instance of each frequency change is considered a ‘deviant tone’, which gradually becomes a ‘standard’ through repetition. Auditory oddball paradigms reveal characteristic early (<300ms) MEG responses to standard and deviant tones. The differential response to these tones (the Mismatch Negativity, MMN) is abnormal in many neurological diseases (Boly et al., 2011; Naatanen et al., 2011; Hughes et al., 2013). The MMN has been proposed to represent a prediction error in hierarchical frontotemporal networks (Garrido et al., 2009b; Phillips et al., 2015). However, earlier models did not reveal the mechanisms of laminar or synaptic function that generate the MMN within the frontal and temporal cortex.

To examine laminar-level dynamics in response to auditory stimuli we used an extended-DCM. In six connected frontotemporal regions (based on Phillips et al., 2015, 2016), we used a conductance-based canonical mean-field cortical modelling scheme (Moran et al., 2013; Marreiros et al., 2015). We introduce cortico-thalamic cells with intrinsic conductances implicated in burst-firing that enable the model to generate beta activity involved in the transfer of deep-layer information (Roopun et al., 2008a, 2010; Bordas et al., 2015; Michalareas et al., 2016). We also employ separate inhibitory interneuronal populations for superficial and deep pyramidal cells (e.g. Jiang *et al.*, 2015). These extensions improve the model’s functionality in terms of cortico-cortical and cortico-thalamocortical transmission and provide a substrate for the greater separation of laminar dynamics. We tested the model’s ability to accurately generate evoked magnetoencephalographic responses (i.e. event related fields, ERF).

We used the drug tiagabine to test how well the neurophysiological model could identify changes in the causes of neuronal dynamics. Tiagabine inhibits re-uptake of the inhibitory neurotransmitter gamma-amino-butyric acid (GABA), which is critical for the generation of physiological responses and rhythms in local and global processing (Whittington et al., 2000). This pharmacological specificity provides a more controlled test of dynamic causal models than autoimmune (Symmonds et al., 2018) and genetic channelopathies (Gilbert et al., 2016).

Using parametric empirical Bayes to optimise the model across participants and drug conditions we examined how GABAergic dynamics in the model are altered by tiagabine. Based on the hypothesis that prediction and prediction error depend on short-term GABAergic plasticity (Castro-Alamancos and Connors, 1996; Garrido et al., 2009a; Mongillo et al., 2018; Spriggs et al., 2018), we predicted that upper and lower hierarchical frontotemporal processing would be differentially affected by tiagabine during standard and deviant tones.

## Materials and Methods

### Experimental Design

We undertook a randomised placebo-controlled double-blind crossover study of the effects of tiagabine in 20 healthy adults (aged 67.5±4.2, ten male). Participants had no neurological or psychiatric illness and were recruited from the MRC Cognition and Brain Sciences and Join Dementia Research volunteer panels. The study was approved by the Cambridge Research Ethics Committee and written informed consent was acquired, in keeping with the declaration of Helsinki.

Neurophysiological responses were measured in an auditory roving oddball paradigm (Garrido et al., 2008). Binaural sinusoidal tones were presented in phase via ear-pieces for 75 ms (with 7.5ms ramp up and down at start and end of the tone), at 500 ms intervals. The frequency of the tone increased or decreased in steps of 50 Hz (range 400 – 800 Hz) after 3 to 10 repetitions. Auditory thresholds were assessed in quiet at 500, 1,000, and 1,500 Hz. Tones were presented at 60dB above the average threshold for a standard population through the earpieces in the MEG.

Each participant attended two MEG sessions with a minimum two weeks interval. They received either 10 mg oral tiagabine or a placebo, in randomised order. Bloods were taken 105 minutes later, immediately prior to MEG data acquisition, to coincide with peak plasma levels and CNS penetration (Nutt et al., 2015).

### Data Acquisition and pre-processing

Magnetoencephalography (MEG) used a 306-channel Vectorview acquisition system (Elekta Neuromag, Helsinki) in a light Elekta Neuromag magnetically-shielded room. This consists of a pair of gradiometers and a magnetometer at each of 102 locations, sampled at 1000 Hz. Vertical and horizontal EOGs tracked eye movements and 5 head-position indicator coils tracked head position. A MEG-Compatible 70 channel EEG cap (Easycap GmbH) using Ag/AgCl electrodes positioned according to the 10-20 system was used concurrently. A 3D digitizer (Fastrak Polhemus Inc., Colchester, VA) was used to record >100 scalp data points, nasion and bilateral pre-auricular fiducials. Subjects also underwent T1-weighted structural magnetic resonance imaging (MPRAGE sequence, TE = 2.9 msTR = 2000 ms, 1.1mm isotropic voxels) using a 3T Siemens PRISMA scanner.

MEG data pre-processing included head position alignment and movement compensation 6 headcoils and employed the temporal extension of Signal Space Separation with MaxFilter v2.2 (Elekta Neuromag). The auto-detection of bad channels was combined with manual input of any channels logged as bad during data acquisition. The Statistical Parametric Mapping toolbox (SPM12) (The Wellcome Trust Centre for Neuroimaging, UCL, UK) was used for further pre-processing and analysis, in conjunction with modified and custom MATLAB scripts (MATLAB 2017a, Mathworks, Natick, MA). Data were Butterworth filtered between 1 and 180 Hz, epoched from −100 ms to 400 ms relative to the auditory stimuli and artefact rejected using EOG, EEG and MEG channel thresholding. Spectral analyses were performed using a multi-taper method. The deviant trial was taken as the 1^st^ trial of a train, regardless of the frequency and the 6^th^ trial of a train was modelled as ‘standard’.

Source reconstruction used a forward model estimated using the single shell cortical mesh from each individual’s T1-weighted MR structural scan. After co-registration using the fiducials and head points, local fields (LFs) for 6 sources of interest were source-reconstructed using SPM “COH” method, a combination of LORETA and minimum norm (Pascual-Marqui et al., 1994; Heers et al., 2016). Sources of interest were (MNI coordinates in parentheses): left auditory cortex (LAud; −42, −22, 7), left superior temporal gyrus (LSTG; −61 −32 8), left inferior frontal gyrus (LIFG; −46 20 8), right auditory cortex (RAud; 46, −14, 8), right superior temporal gyrus (RSTG; 59 −25 8) and right inferior frontal gyrus (RIFG; 46 20 8). To create images of induced power, SPM-LORETA was used for source localization of a 5 mm^3^ regular grid at the MMN (150 – 250 ms) time window (100ms in width, regularization=0.05).

Correlation coefficients for comparing the actual and predicted ERFs were calculated using the corrcoef function (Pearson correlation) in MATLAB 2017a for each individual, condition and node. Time-frequency analysis was performed in SPM12 using a multi-taper method with 100 ms windows overlapped by 5 ms and a bandwidth of 3. Frequency bands were split into alpha (8 – 13 Hz), beta (14 – 29 Hz), low gamma (30 – 48 Hz) and high gamma (52 – 80 Hz).

### Neuronal Modelling: an extended canonical microcircuit model

We used conductance-based canonical mean field (CMM) models for evoked responses (Kiebel et al., 2008) utilising canonical microcircuit models (SPM12, DCM10). This approach to neurophysiologically informed modelling using DCM goes beyond descriptive biomarkers by providing a mechanistic link to realistic microscopic processes. A common approach in DCM is to invert the neuronal and spatial forward model as a single generative model, to solve the source reconstruction and biophysical modelling problems jointly by fitting the DCM to sensor data.

However, we modelled source specific responses to suppress conditional dependencies between the neuronal parameters and the parameters of a spatial forward model. This affords more efficient estimators of neuronal parameters, providing the source reconstruction is sufficiently precise given the spatial topography of the network of interest. This has the advantage of compatibility with multiple studies of this task (Muthukumaraswamy et al., 2015; Gilbert and Moran, 2016; Shaw et al., 2017, 2018), including MEG and electrocorticography studies; the chosen network was based on the published bilateral A1, STG, IFG networks associated with the generation of the MMN response. Since this spatial element of the inverse problem was constrained, it is computationally more appropriate to source localise using SPM with prior expected sources. The subsequent DCM was then run on these virtual electrodes.

The DCM included a homologous conductance-based neural-mass model at each of the six anatomical locations, as shown in Figure 1. They comprised 6 cell modules: a superficial pyramidal module (sp), a deep cortico-cortical pyramidal module (dp), a thalamic-projection pyramidal module (tp), a granular stellate module (ss) and separate supragranular and infragranular interneuron populations (si & di). Excitatory autapses existed for all excitatory cell modules and all modules were also governed by an inhibitory self-gain function that provided tonic inhibition to each module. The intrinsic connectivities are shown in Fig. 1a: note the excitatory conductances based on AMPA and NMDA and inhibitory GABA-A and GABA-B conductances. The model is an extension of the SPM conductance-based CMM model (SPM12, 2013): inclusion of separate supra- and infra-granular interneuron populations creates a more biophysically realistic model that allows a greater flexibility of independence of deep and superficial activity than in previous work (Bhatt et al., 2016; Shaw et al., 2018; Spriggs et al., 2018). Additionally, the new ‘tp’ population expressed a hyperpolarization-activated cation current (H-current) and a non-inactivating potassium current (M-current) to provide surrogate intrinsic dynamics involved in the characteristic bursting behaviour of these cells. This, coupled with a different cell capacitance, differentiated the intrinsic activation of the ‘tp’ population from the ‘dp’ population. The populations also differed in their extrinsic connectivities, with ‘dp’ populations forming cortico-cortical connections and ‘tp’ populations allowing for cortico-thalamocortical connections. Thalamic activity was not specifically modelled but is represented by an 80 ms delay in connectivity.

**Figure 1.**
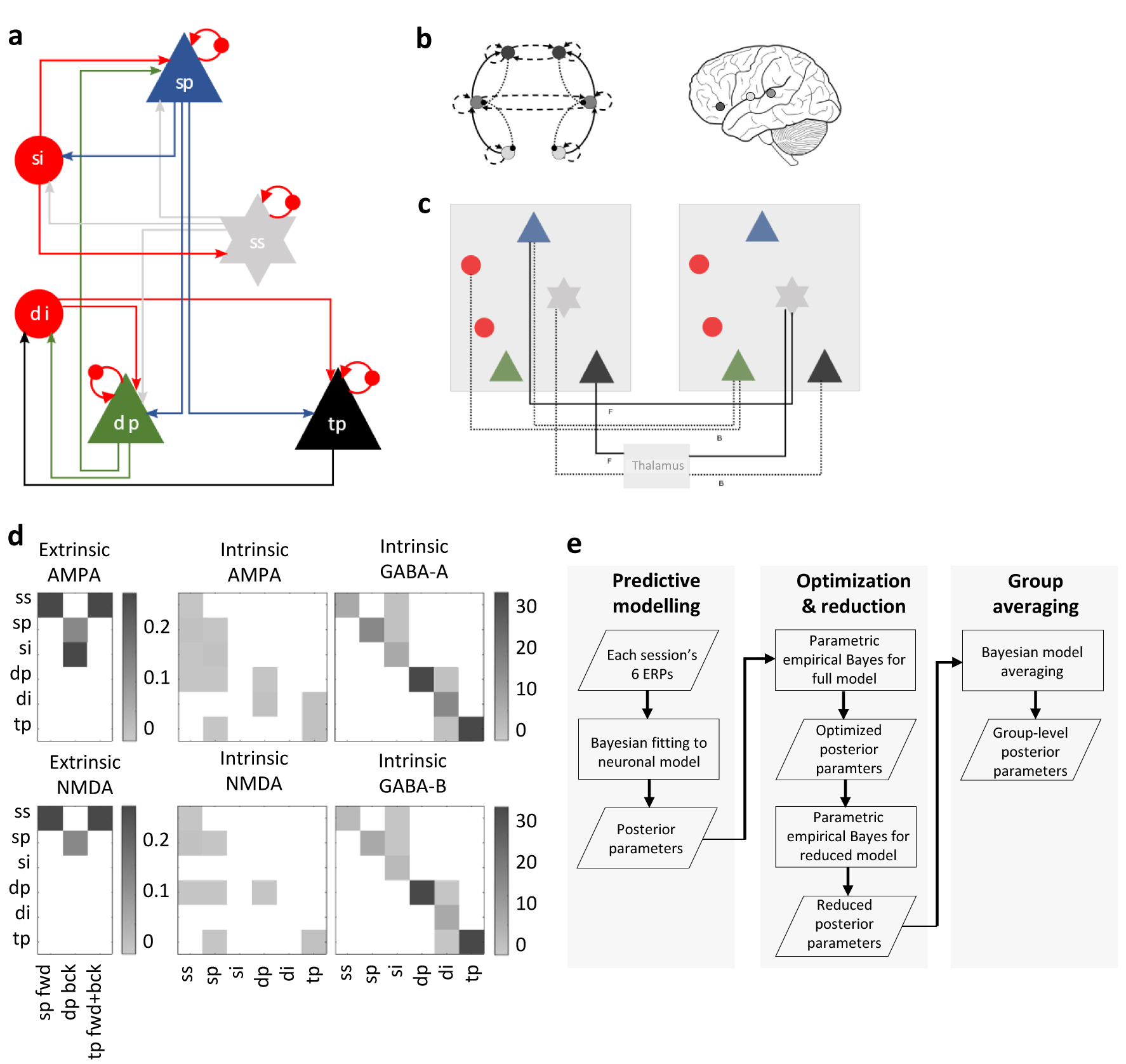
The neuronal model. a. Intrinsic connectivities found in all nodes between layer 4 stellates (ss), inhibitory interneurons (ii), superficial pyramidal modules (sp) and deep pyramidal modules (dp).
b. All 6 nodes used are represented as a network on the left, showing the extrinsic connectivities (solid line = forward; dotted line = backward; dashed line = lateral). A left hemisphere representation of these bilateral nodes in primary auditory cortex, superior temporal gyrus and inferior-frontal gyrus (light, medium and dark grey, respectively).
c. A detailed view of the extrinsic population connections for forward (solid lines) and backward (dotted lines) connections.
d. Matrices of the extrinsic and intrinsic connectivity weights, all of which had a permitted variance of 1/16.
e. A process flow describing the steps taken in the meta-analysis phase.

Extrinsic connectivity between the six nodes is shown in Fig. 1b, with the detailed extrinsic population connections shown in Fig. 1c. In keeping with the established principle of differential cortical laminar projections of feed-forwards vs feedback connectivity (Bastos et al., 2012), backward connections are facilitated by the ‘dp’ cells terminating on ‘sp’ and ‘si’ cells, whilst forward connections run from ‘sp’ cells to ‘ss’ cells. Cortico-thalamo-cortical connections originate from ‘tp’ cells and terminate following a thalamic delay at layer 4 ‘ss’ cells. The presence or absence of connections between nodes was based on the fully connected models from Phillips et al., (2015) and Shaw et al., (2019), which in turn were derived from Garrido et al., (2008). This was used for the basis of an iterative process to find the most likely reduced model (described below).

A Gaussian kernel (peak 60 ms, half-width 8 ms) represented auditory input to layer 4 stellates in bilateral auditory and inferior frontal cortex.

### Bayesian Modelling and Statistical Analysis

We used Bayesian model inversion and selection to identify the best explanation for subject-specific data, in terms of neuronal and biophysical parameters. Parametric Empirical Bayes was used for group inferences and to examine drug effects.

The DCM was inverted to source-reconstructed ERF data for the 6 nodes for each subject. Data were filtered between 0–48 Hz and a Tukey window was applied that did not attenuate signals 50 ms before or 350 ms after stimuli. Model inversion was run separately for the standard and deviant trials and passed to second level Parametric Empirical Bayesian with contrasts for both trial types and drug conditions. All intrinsic and extrinsic AMPA, NMDA and GABA-A conductance scalings could vary independently in a manner that assumed symmetry between the two hemispheres. The prior means and permitted variances are summarised in Table 1.

**Table 1.** Model parameters. Parameter values used by the neuronal model are shown with their permitted variances.

| Parameter grouping | Parameter | Initial value | Permitted variance |
| --- | --- | --- | --- |
| Decay<br>Constants, $\tau$ (ms) | AMPA $\tau$ | 4 | 1/16 |
| | NMDA $\tau$ | 100 | 1/16 |
| | GABAA $\tau$ | 16 | 1/8 |
| | GABAB $\tau$ | 200 | 1/8 |
| | $I_M \tau$ | 160 | 0 |
| | $I_H \tau$ | 100 | 0 |
| Misc. strengths | K<sup>+</sup> leak G | 1 | 0 |
|  | Background V | 2.17 | 1/32 |
| Reversal potentials (mV) | Na<sup>2+</sup> reversal | 60 | 0 |
|  | Ca<sup>2+</sup> reversal | 10 | 0 |
|  | Cl<sup>-</sup> reversal | -90 | 0 |
|  | K<sup>+</sup> reversal | -70 | 0 |
| | $I_H$ reversal | -100 | 0 |
| Firing threshold (mV) | $V_T$ (all pops) | -40 | 0 |
| Firing precision | $V_X$ (all pops) | 1 | 1/32 |
| $I_H$ I-V slope | $V_{HX}$ | 300 | 0 |
| Cell<br>Capacitances (pF) | $ss_C$ | 200 | 1/32 |
| | $sp_C$ | 150 | 1/32 |
| | $si_C$ | 50 | 1/32 |
| | $dp_C$ | 400 | 1/32 |
| | $di_C$ | 50 | 1/32 |
| | $tp_C$ | 200 | 1/32 |
| Delays (ms) | intrinsic | 2 | 1/32 |
|  | extrinsic<br>cortico-cortical | 16 | 1/32 |
|  | extrinsic thalamo-cortical | 80 | 1/32 |

Variational Bayesian statistics using the Laplace approximation determined the probable parameter space given the neuronal model and the data (Friston et al., 2007). The full model parameter space was reduced by iteratively searching for dependencies in this parameter space and systematically removing parameters not contributing to the free energy of the system (Henson et al., 2011). The optimised reduced model comprises all those parameters and connections found to contribute significantly to the system temporal dynamics. The parameter distributions from this reduced model were used to create a Bayesian average model of parameters that differ significantly across the contrasts of trial types and drug conditions. The process flow is summarised in Fig 1e.

Frequentist statistical methods quoted in the main text used MATLAB (2017a, Mathworks, Natick, MA). Classification of data into the placebo and drug conditions used a linear SVM in the Classification Learner Application in MATLAB 2017a (Mathworks, Natick, MA) with 5-fold cross-validation approach.

*Code Accessibility:* The custom neuronal model used to generate these results is available at [**address on acceptance**] and works in conjunction with SPM12.

## Results

### Event related fields and induced spectral power

Event related responses to standard and deviant trials were in line with previous findings (Hughes and Rowe, 2013; Phillips et al., 2015, 2016) (Fig. 2a, first and second rows) and show the expected M100, the primary response after the onset of a tone (80-120 ms), a difference signal (MMN) between the standard and deviant trials (150-250 ms) and an M300 visible in frontal nodes (250-380 ms). The M100 was significantly reduced by tiagabine on standard and deviant trials, in left temporal nodes (A1, and STG p<0.05, paired t-test), whereas the later response leading into the M300 was significantly reduced only on deviant trials in L/R IFG (p<0.05).

**Figure 2.**
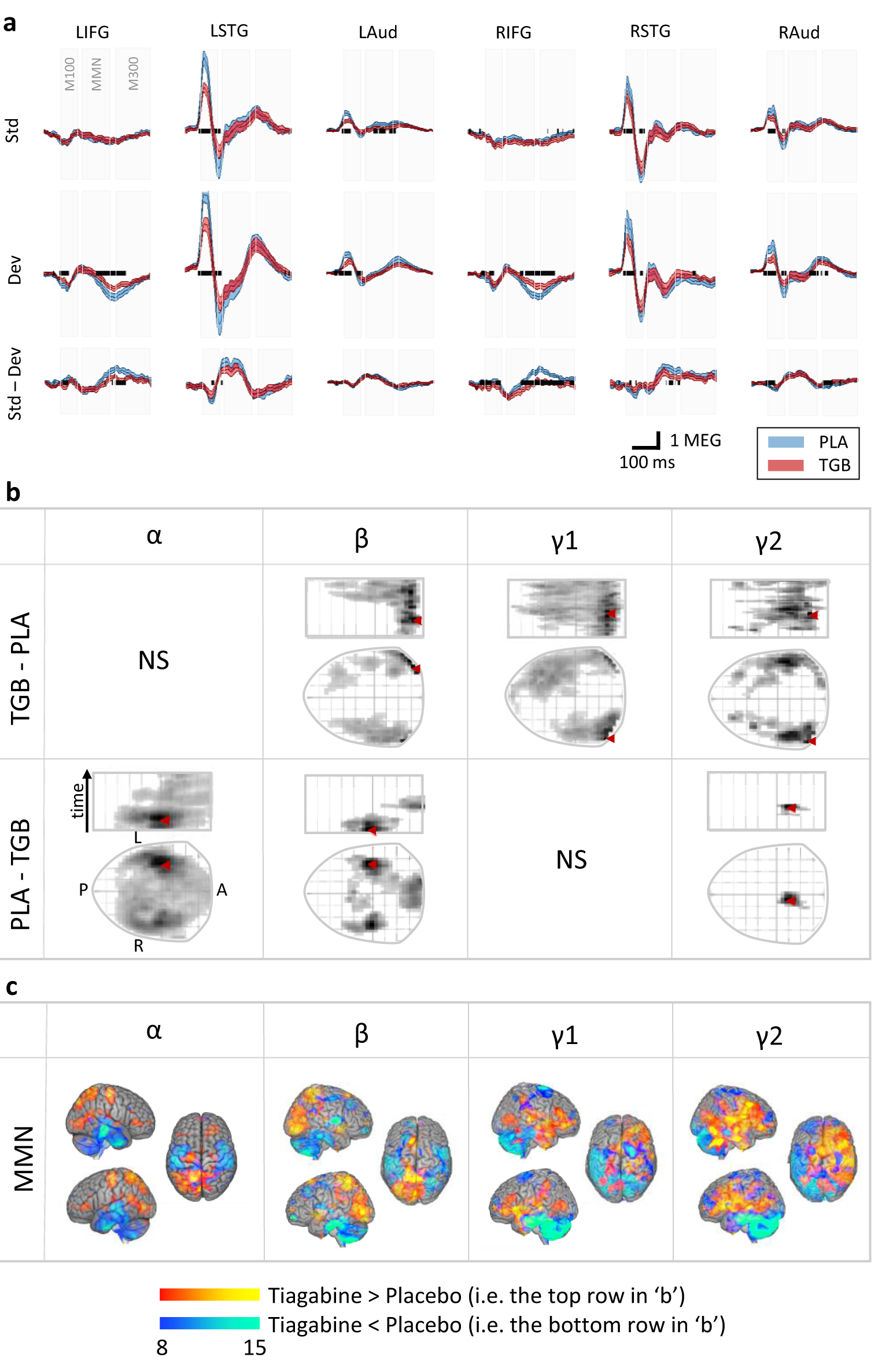
Event Related Fields (ERFs) a. Mean ERFs across all subjects for all six nodes for the standard and deviant trials from 0-380ms. The difference wave (MMN) is also shown. ERFs from the placebo condition are shown in blue and from the tiagabine condition in red. Significant (p<0.05) changes with time across the drug condition are shown as a thick black line within each axis. Shaded areas represent the standard error (SEM).
b. Significant differences for induced spectra power were found in the alpha (α), beta (β) and lower and higher gamma bands (γ1 and γ2). Here they are shown as flat scalp maps (lower plots) with rostro-caudal activity versus time (upper plots). The time axis runs from 0–380 ms post-stimulus.
c. Source-reconstructed T-contrasts created for those frequency bands showing significant spatial changes across the drug condition in the 135 – 235 ms time window.

The difference waveform (i.e. the deviant – the standard) reveals a typical biphasic MMN between 150-250ms, observed in primary auditory cortex and STG (Fig. 2a, third row). Tiagabine significantly reduced the second peak of the MMN (p<0.05) with bilateral IFG nodes and RSTG showing reductions in the first peak of the mismatch response on tiagabine (p<0.05). As with the deviant response, LIFG showed a significant reduction of the later MMN peak and the M300 on tiagabine (p<0.05).

The temporal profile of spectral power differences (see Methods for time-frequency analysis) matched that of the ERFs, including spectral counterparts to M100, MMN, continuing through the M300 window (Fig. 2b&c). During the M100, alpha-power (8-12 Hz) decreases on tiagabine were localized to temporal cortex and beta (14-29 Hz) decreases more prominently to posterior temporal cortex. During the MMN, increases in low and high gamma (30-48 Hz and 52-80 Hz respectively) were observed broadly across right frontal cortex, including IFG. Low gamma also showed increases in right temporal cortex.

Such changes in the observed spatiotemporal physiology on tiagabine will be dependent on changes in local and global network connectivity. The extended conductance-based dynamic causal model was therefore used to infer the causes of the observed physiological changes.

### The Dynamical Causal Model

The modelled responses are explained in terms of the parameters of the optimised model. Using parametric empirical Bayes, model parameters were compared across the standard and deviant conditions, as well as across the placebo and tiagabine conditions. Figure 4 shows the effect of tiagabine on the intrinsic GABAergic connectivity, assuming symmetry (three bilateral averaged nodes are shown). We confirmed that tiagabine significantly increases tonic GABAergic inhibition (posterior probability given for each parameter in Fig. 4a). This was seen primarily in the deep layer pyramidal and interneuron populations (Fig 4a).

**Figure 3.**
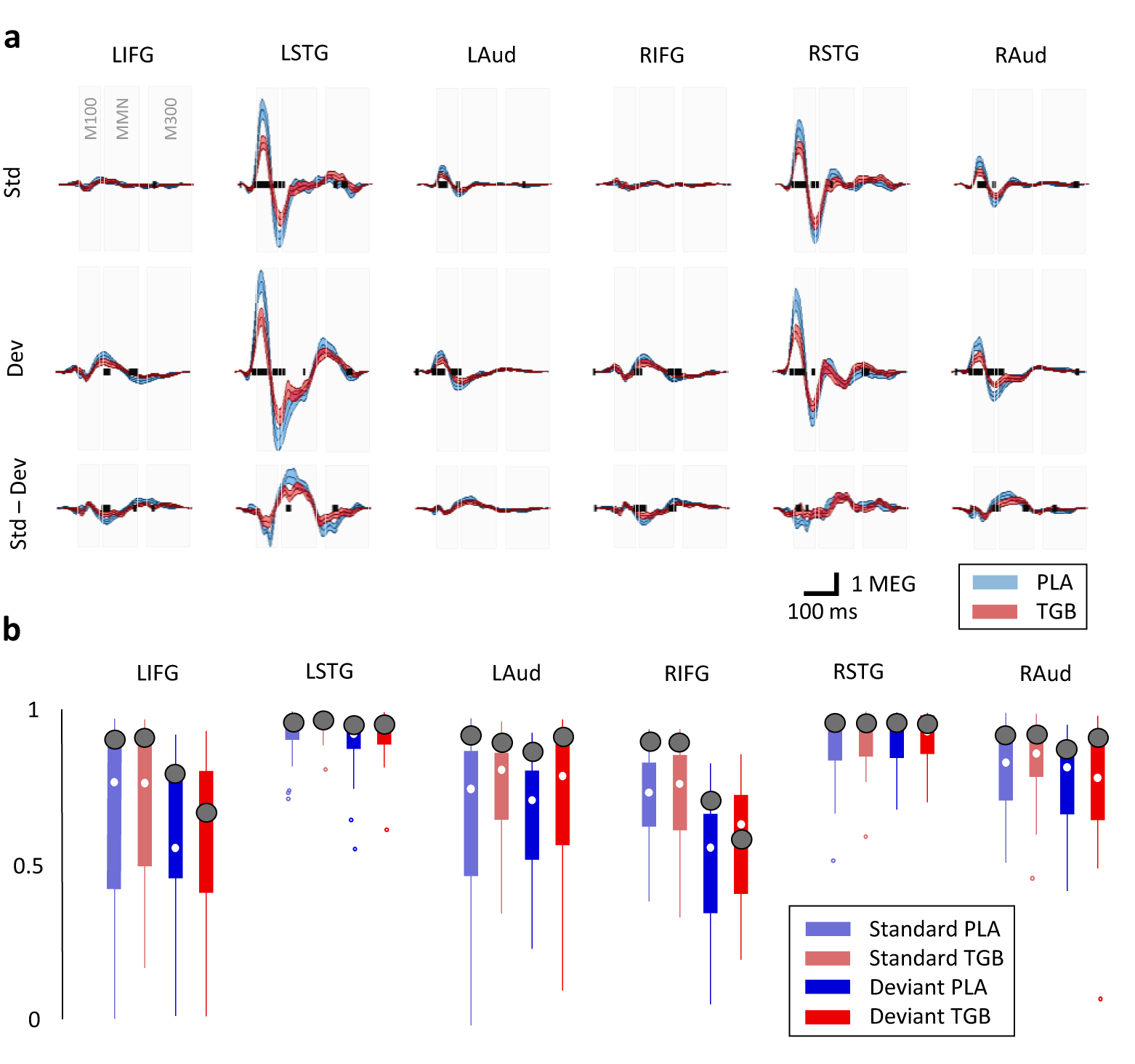
Comparison between model and data. a. Predicted ERFs are shown for the standard and deviant conditions, along with the difference wave (Std–Dev). The placebo and tiagabine conditions are depicted in blue and red respectively with significant differences (p<0.05) shown as a thick black line within each axis.
b. Correlation coefficient between prediction and data for each node and each condition. Boxplots represent the distribution over subjects with small dots representing outliers and larger black circles representing the meaned response of all subjects.

**Figure 4.**
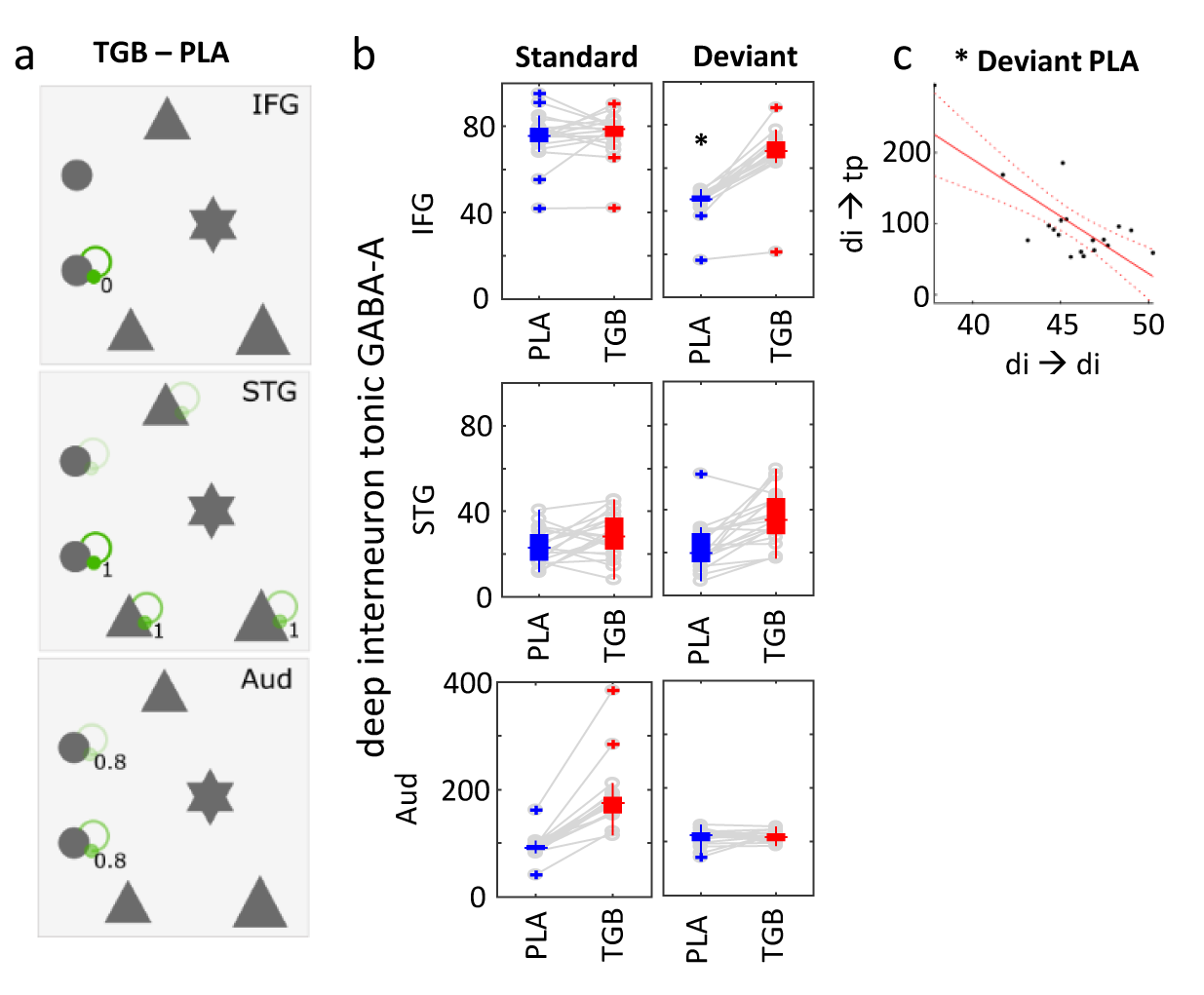
Prediction of hidden states. a. Significant differences in the modulation of GABA-A synaptic scaling for each of the three symmetric nodes. Green/red show significantly greater/lesser GABA-A synaptic scaling for tiagabine than the placebo.
b. Tonic GABA-A scaling on deep interneurons in IFG, STG and Aud, for each individual, plotted for the placebo and tiagabine conditions. The standard and deviant conditions are plotted separately in the left and right columns respectively.
c. Linear fit with 95% confidence bounds for tonic GABA-A scaling on deep inhibitory neurons vs phasic GABA-A scaling from deep inhibitory neurons to thalamic projecting pyramidals (p=1.7e-4, significant to p<0.01, Bonferroni corrected).

In keeping with the functional differentiation of upper versus lower levels in a hierarchical neural network with backwards-prediction and forward-prediction error, there was an interaction between the effects of tiagabine and condition between regions: Fig 4b compares GABA-A conductance scaling on deep interneurons between placebo and tiagabine conditions, plotted for each individual.

The differences were significant (standard paired t-test, p=1.1e-8) between the two groups in primary auditory areas for the standard condition, and in IFG for the deviant condition.

The correlation between tonic and phasic inhibition was also explored for each region and condition and a strong negative relationship was found between the tonic inhibition of deep inhibitory cells and their phasic inhibition onto cortico-thalamic cells (Fig. 4c p=9.0e-8, Bonferroni corrected).

Finally, we used the estimated parameters for the 20 individuals in a linear support vector machine with 5-fold cross-validation to classify the conditions under which data were acquired and for which the models were therefore optimised. Parameter based classification reached 92.5% accuracy.

### Discussion

The principle insights from this study are (i): an extended conductance-based canonical mean-field method of dynamic causal modelling (“ext-DCM”) is tractable and accurate for generating event-related fields that match those observed by magnetoencephalography; and (ii) we confirmed the modulation of GABAergic dynamics by the GABA-reuptake inhibitor tiagabine, opening the way for psychopharmacological studies in health and disease with the mechanistic precision afforded by using ext-DCMs as generative models.

We demonstrate that key features of the intrinsic connectivity within-regions changes across conditions in simple MMN paradigm, but they are of generalised relevance to hierarchical network models of cognition such as speech (Cope et al., 2018), semantic (Adams et al., 2019) and visual perception (Muthukumaraswamy *et al.*, 2013). Moreover, the laminar and pharmacological specificity provided by the ext-DCM has the potential to quantify neuropathology in dementia, developmental and psychiatric disorders (Duyckaerts et al., 1986; Kinoshita et al., 1996; Ferrer, 1999; Ji et al., 2018; Shaw et al., 2018).

In the following sections, we first discuss how MEG quantifies the effects of tiagabine on cortical dynamics. We then consider additional insights from biophysically informed DCMs of hierarchical brain networks, illustrating mechanistic explanations of the observed population dynamics.

### Understanding the MMN in terms of short-term plasticity

The drug modulation of GABA resulted in complex dynamics across the trial types, implicating both local tonic-phasic effects and global hierarchical effects. Repetitive activation with the same stimulus results in a dampening effect on the ERF (reduction in N1/N2 by 6^th^ repetition), Fig 2. We predicted higher tonic inhibition in the deep layers during the repetition state – namely, increased tonic inhibition in cortico-cortical circuitry in prefrontal cortex and cortico-thalamo-cortical circuitry in temporal cortex – which we interpret as local short-term plastic changes in deep-layer inhibition (Knott et al., 2002; Hensch, 2005; Jääskeläinen et al., 2007) that regulates salient information (Mongillo et al., 2018).

The model confirmed that the effect of tiagabine was to increase extracellular GABA concentrations with a marked increase in tonic inhibition, associated with overspill of GABA onto extra-synaptic receptors (Semyanov et al., 2004). The effect was modulated differently in primary and secondary processing areas: for tonic inhibition of deep interneurons the drug’s efficacy was highest in prefrontal cortex for deviant trials and in auditory cortex for standard trials. Figure 4b shows that whereas a drop in deep interneuron tonic inhibition was observed on deviant trials, tiagabine abolished the effect. We speculate that the drop in tonic inhibition at the presentation of a deviant tone relates to homeostatic competition between phasic and tonic inhibition (Wu et al., 2013), with phasic activation of deep-layer projections being necessary for feedback of top-down information on context. Increasing tonic inhibition likely decreases the interneuron population activation (Semyanov et al., 2004), leading to decreased phasic inhibition onto deep pyramidal cells. This relationship was confirmed and is shown in Fig 4c between tonic inhibition of deep IFG interneurons and phasic inhibition of deep IFG thalamic-projection neurons.

### GABA-ergic modulation of evoked and induced responses

Tiagabine has a range of effects on oscillatory dynamics, in beta and gamma ranges, which may influence behaviour (Coenen et al., 1995; Magazzini et al., 2016; Port et al., 2017; Wyss et al., 2017). It remains a challenge to relate systemic drug effects with such local frequency-spectral phenomena. However, it has been proposed that beta-band activity is associated with infragranular cortical projection neurons with intrinsically bursting profiles (Groh et al., 2010; Roopun et al., 2010; Kim et al., 2015). Here we found that, on tiagabine, induced beta-band activity was reduced in temporal areas. This relates to the model prediction that tonic inhibition is increased on intrinsically bursting thalamic projection neurons in STG, and not phasic inhibition, which could increase rebound bursting via intrinsic M- and H-currents (Roopun *et al.*, 2008; Roopun *et al.*, 2008b).

Conversely, it has been shown that gamma-band activity is dependent on the GABA-A receptor activation and the phasic interplay of interneuron-pyramidal cell networks, particularly in the superficial layers (Buffalo et al., 2011; Whittington et al., 2011). Our evidence from the mismatch temporal window (Fig. 2b) indicates peak gamma increases occurring at the start of the mismatch period. This is consistent with thalamic input (Di and Barth, 1992, 1993; Sukov and Barth, 2001) leading to an envelope of gamma activity in the superficial layers of cortex during audition (Metherate and Cruikshank, 1999).

Overall, the observed dynamics and the model posterior parameters are consistent with our knowledge of network activation within the context of beta- and gamma-rhythm generation in cortex and how increases in endogenous GABA could manifest.

### Generative models of drug effects on cognitive physiology

The drug’s effect was largely confined to deep cells that in turn connect to superficial cells, with tiagabine reducing deep-layer influences on superficial layers. As we modelled evoked activity it is difficult to speculate on how this influences gamma activity across the network, however a reduction in deep-layer influence may increase local cortical processing associated with gamma-band activity in the superficial layers. Under the assumption that their GABA levels are lower in older versus younger adults, tiagabine acts restoratively to increase gamma-band activity by altering the balance of activity across layers. This is corroborated with lower frequency band activity, dependent on GABA (Mathias et al., 2001). Finally, we speculate that the reduced M100 seen on tiagabine is a consequence of the widespread increased tonic inhibition predicted by the model (Fig. 4), causing a general reduction in local population activity.

### Study limitations

Our study was motivated by the need for mechanistic studies of human cortical function, underlying cognition, disease and therapeutics. Despite support for our three principal hypotheses, and background validation studies (Moran et al., 2014), evidence from one study may not generalise to other tasks and populations. There are some study-specific considerations that limit our inferences, in relation to our participants, our model, and drug of choice. For example, our participants were healthy, but they were older than those studied by Nutt et al (2015), and therefore have normal age related changes in GABA (Gao et al., 2013; Eavri et al., 2018), that could interact with the effects of tiagabine (Nutt et al., 2015).

Our neuronal model provides a simplified substrate for the neurophysiological processes. It is more detailed than previously canonical microcircuit convolution models (Moran et al., 2013), in an effort to improve the modelling of specific dynamics in the form of cell populations, their differing connectivities, synaptic time constants and voltage-gated conductances relevant to this cortical micro-circuitry and task. The extended model can produce a wide spectrum of oscillatory responses, including both superficial gamma rhythms and deep beta rhythms (Roopun et al., 2006; Kramer et al., 2008; Whittington et al., 2011). It can incorporate delayed activity associated with local, cortico-cortical and cortico-thalamo-cortical connections. Currently, this system is a simplified network acting as a neural mass, and as such it can represent relevant cortical interactions involved in ERF generation in the context of this task and study. It does this by allowing forward and backward modulation of activity between deep and superficial layers, where synaptic time constants corroborate with standard GABA, NMDA and AMPA receptor decays. The six specified nodes are commonly cited in the literature in the context of this task (Garrido et al., 2009b; Phillips et al., 2015). Although they are not a complete representation of possible network configurations, they have nevertheless been shown to capture critical aspects of cortical function: here the network has been supplemented with modelled exogenous and endogenous inputs via thalamus. Where parameters derived from DCMs are used for frequentist statistical tests, they have excellent reliability across sessions and sites, and similar power to fMRI and EEG studies (Rowe et al., 2010; Goulden et al., 2012; Bernal-Casas et al., 2013). However, such a frequentist approach is obviated by the direct inferences on posterior probability inherent in the Bayesian inference of DCM, and the use of Parametric Empirical Bayes in particular for group studies.

tiagabine is a relatively specific blocker of GAT-1 at the concentrations used, but does not distinguish between the mechanisms activated by GABA (Bowery et al., 1987; Mody and Pearce, 2004; Lee and Maguire, 2014). The timing of the magnetoencephalography coincided with expected peak plasma levels, but levels may vary between individuals and future studies could in principle include levels as a covariate of interest, or model time-varying responses in relation to drug levels (Muthukumaraswamy et al., 2013b).

In conclusion, we have used a conductance-based model of cortical neuronal dynamics to study GABA-ergic interactions and probe laminar-specific physiological responses to tiagabine. The model accurately generated physiological data that matched the MEG responses and confirmed the effect of tiagabine on tonic GABA-A inhibitory gain within frontal and temporal cortical circuits. Our data provide support for mechanistic studies of neurological disorders, including but not limited to GABAergic impairments (Murley and Rowe, 2018). They also point to new approaches for experimental medicine studies in humans that aim for the laminar, cellular or synaptic precision made possible in new generations of dynamic causal models.

## Acknowledgements

This work was funded by the Wellcome Trust (103838), the National Institute for Health Research Cambridge Biomedical Research Centre and the Medical Research Council (MC_U105597119 & MC_U_00005/12) and Holt Fellowship. We thank the PSP Association & FTD Support Group for raising awareness of the study.

